## Supplementary data for "Tumor-infiltrating nerves functionally alter brain circuits and modulate behavior in a mouse model of head-and-neck cancer"

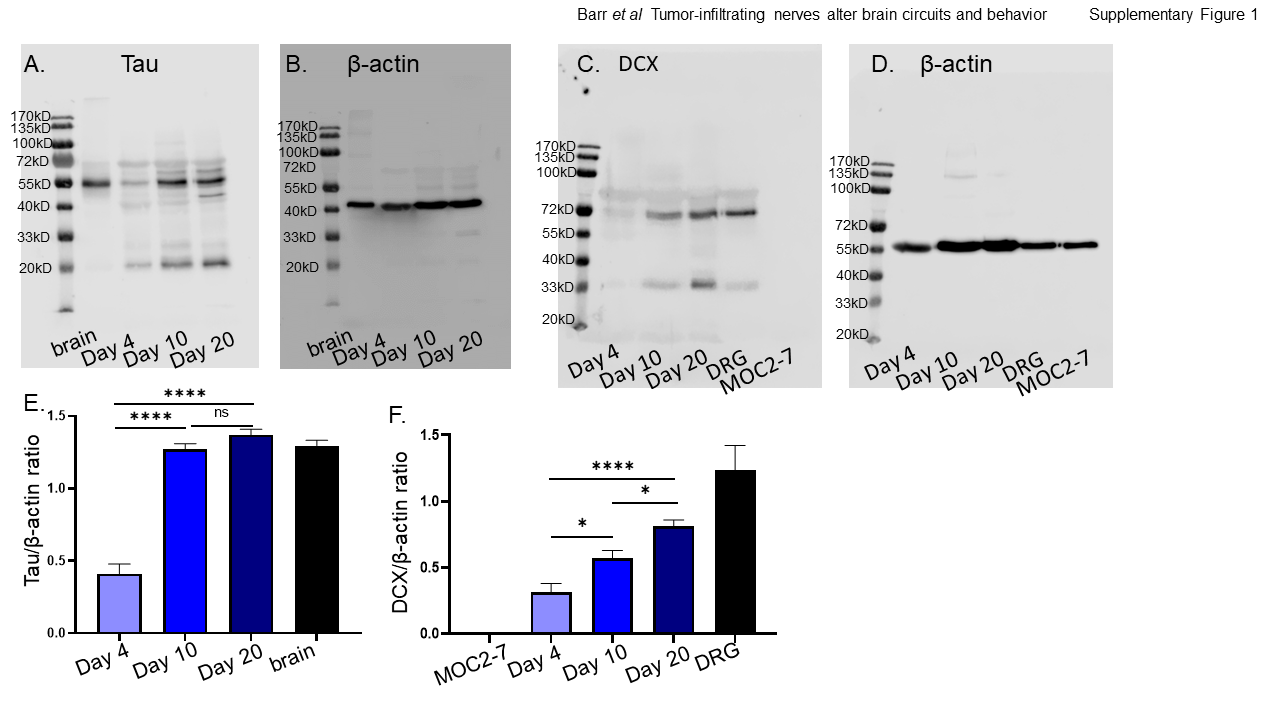

**Supplementary Figure 1. Full westerns for tumor innervation time course.** Whole tumor MOC2-7 lysates were harvested at the indicated days post-tumor innervation and analyzed by western blot for Tau (**A**), β-actin (**B**) (loading control for panel A), (**C**) Doublecortin (DCX), (**D**) β-actin (loading control for panel C). Densitometric quantification of western blots in panels A and B (**E**) and panels C and D (**F**). Statistical analysis by one-way ANOVA with post-hoc Tukey test; n=3 mice/time point, n=4 technical replicates. *, p<0.05; ****, p<0.001

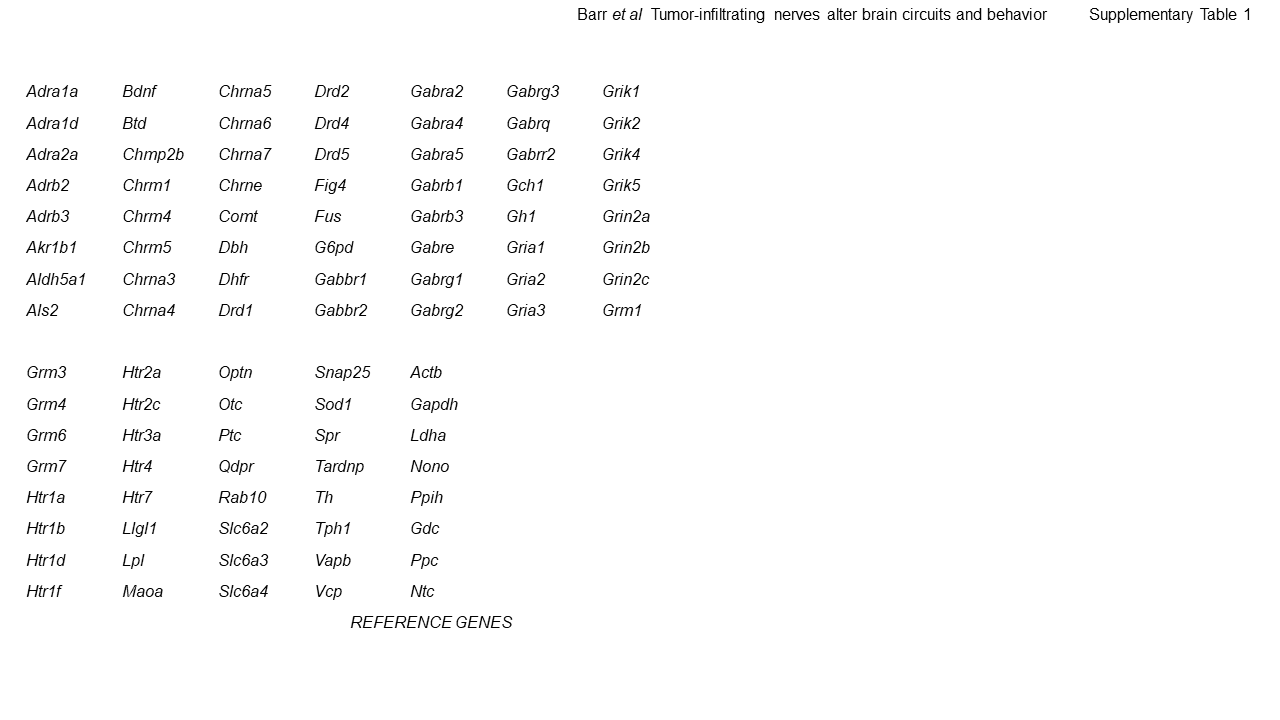

**Supplementary Table 1.** List of mouse genes assayed on the ScienCell Gene Query Neuronal Transmission and Membrane Genes plate.

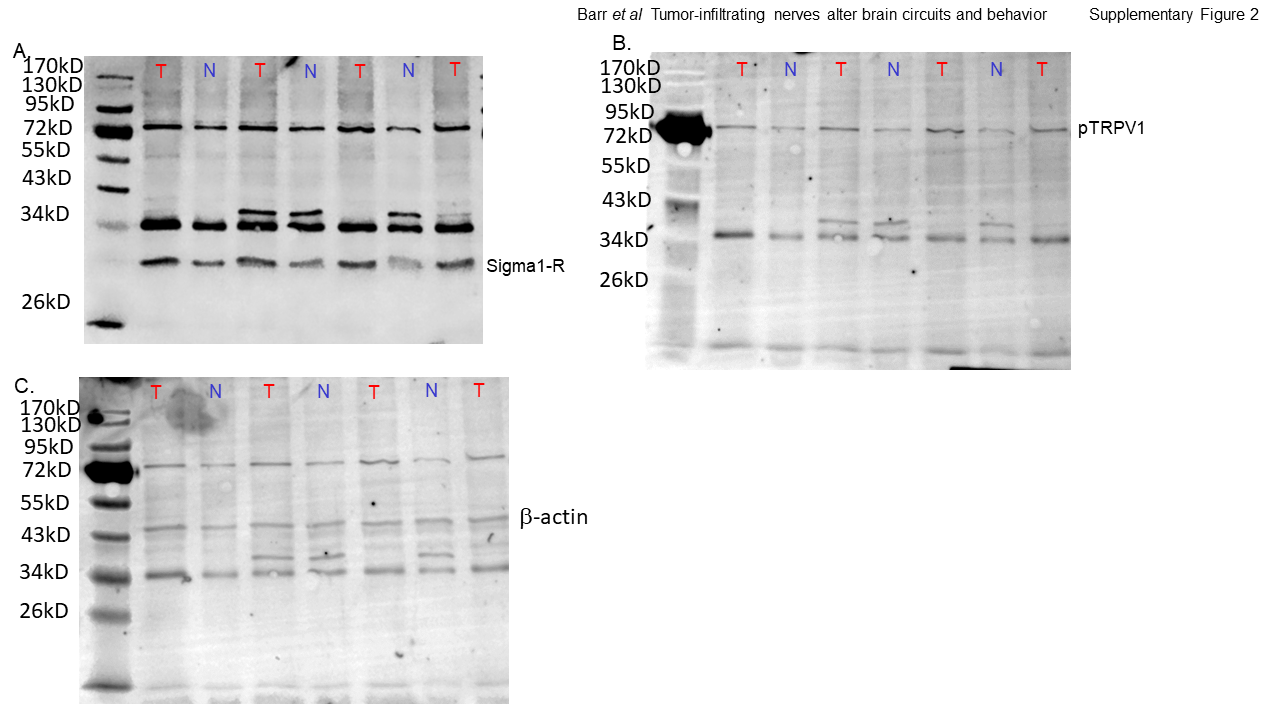

**Supplementary Figure 2. Full westerns.** Full western blot of trigeminal ganglia (TGM) from MOC2-7 tumor bearing (T) or non-tumor bearing (N) mice blotted for the sigma-1 receptor (**A**), phosphorylated TRPV1 (pTRPV1) (**B**) or β-actin (loading control) (**C**). Full western blot from panel A re-probed for blots (**B**) and (**C**) were not stripped in between probing for different proteins.

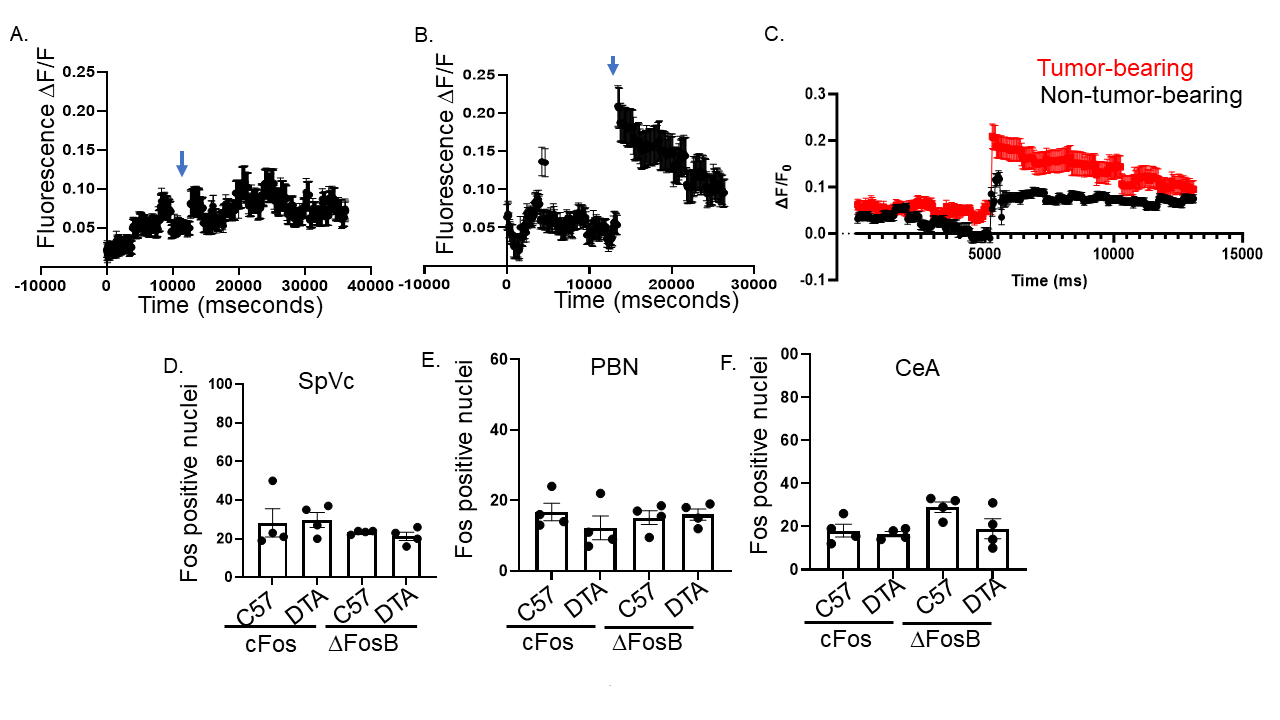

**Supplementary Figure S3.** Representative calcium recordings for a single neurons from a brain slice from control (**A**) or tumor-bearing (**B**) animals. Arrow denotes time of KCl stimulation. Curve of change in Ca+2 fluorescence ex viso brain slices harvested from mice with oral tumors and controls without tumor (n=3 brains/group and n=3 slices analyzed/brain) (**C**). Quantification of cFos and ΔFosB positively stained neurons in the spinal nucleus of the trigeminal (SpVc)(**D**), parabrachial nucleus (PBN) (**E**) and central amygdala (CeA) (**F**) from control C57Bl/6 (C57) or TRPV1^cre^::DTA^fl/wt^ (DTA) animals. No significant differences found.

A.	B.

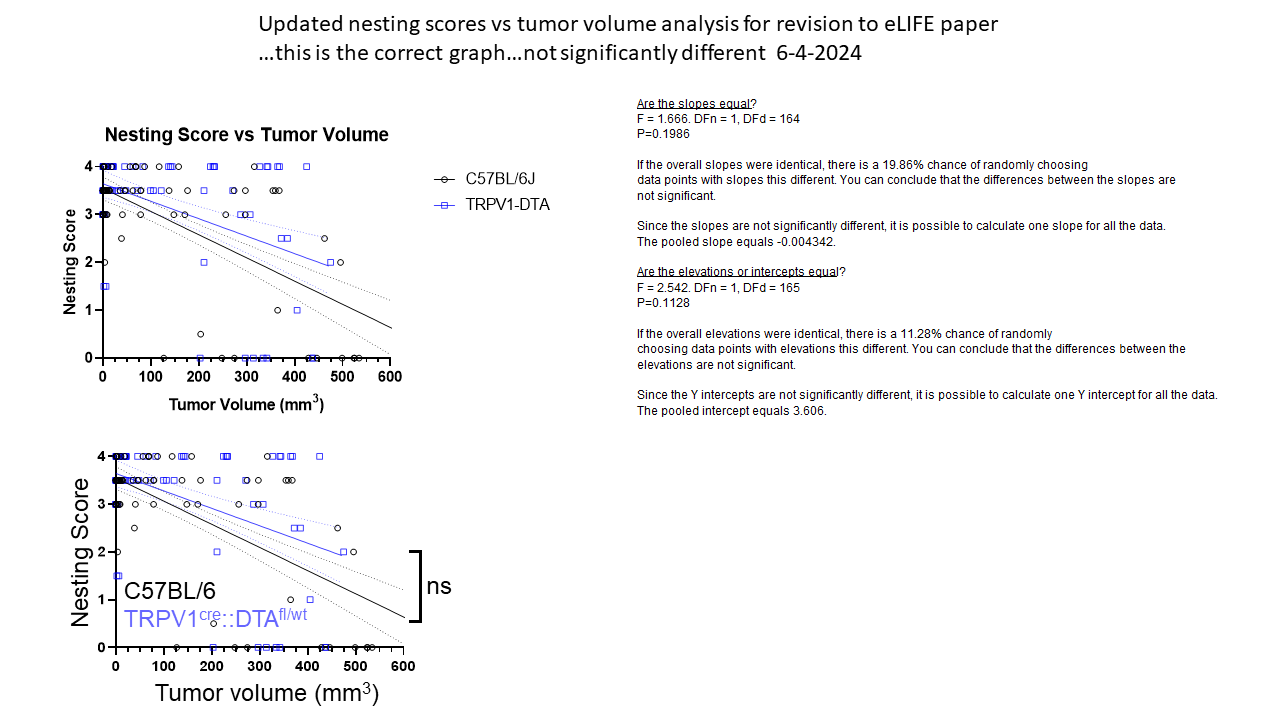

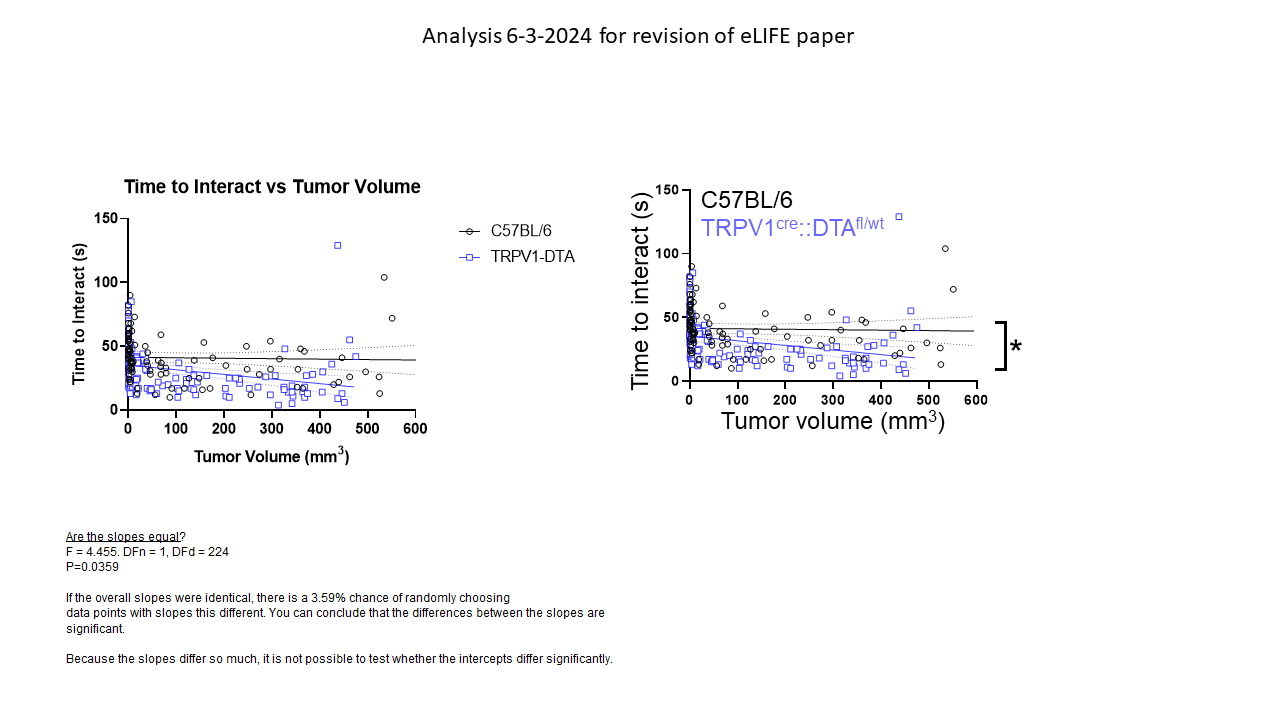

**Suppl Fig 4. Tumor volume does not account for nesting differences.** (A) Nesting data from MOC2-7 oral tumor-bearing C57BL/6 (black, n=15 mice) and TRPV1^cre^::DTA^fl/wt^ (blue, n=14 mice) were graphed as a function of tumor volume. A simple linear regression analysis was completed to determine best fit values. The slopes of the lines are not significantly different. ns, p=0.1986. F=1.666; DFn=1; DFd=164. (B) These same mice underwent the cookie test and the times to interact with the cookie similarly graphed as a function of tumor volume. A simple linear regression analysis was completed to determine best fit values. *, p= 0.0359, F= 4.4459, DFn= 1, DFd= 224.

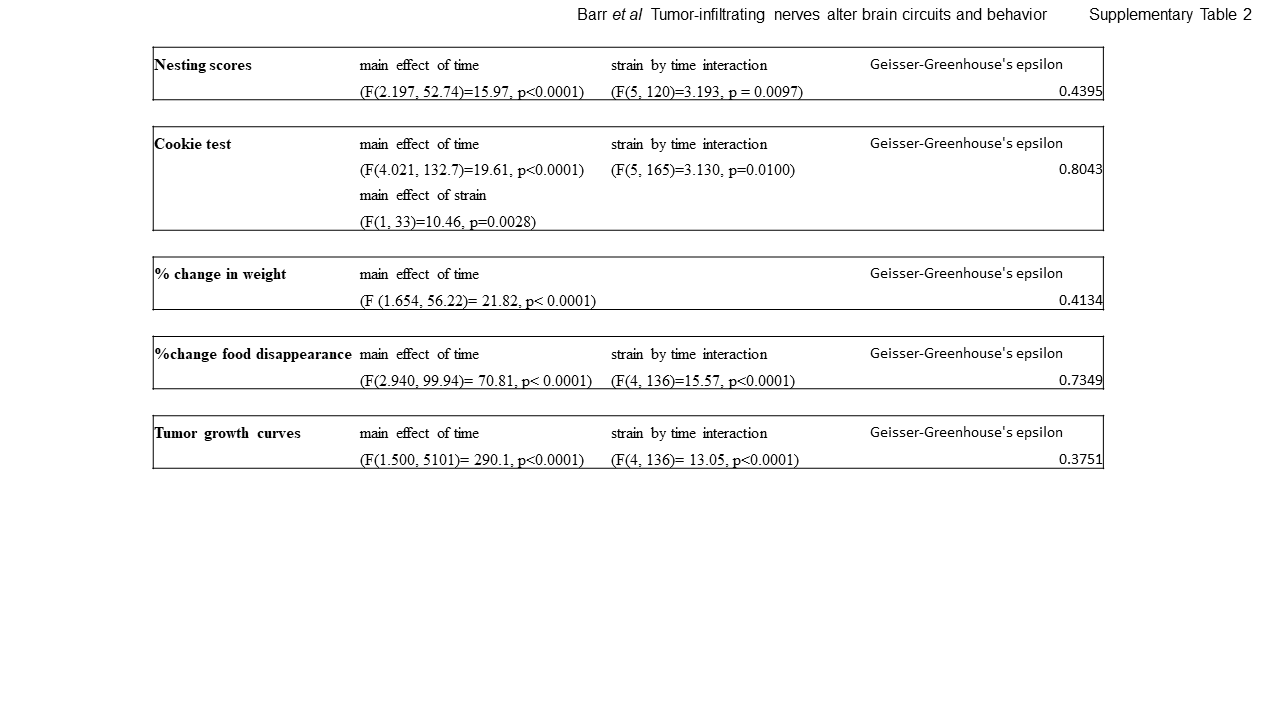

**Supplementary Table 2.** Significant effects for the statistics presented in Figure 4.

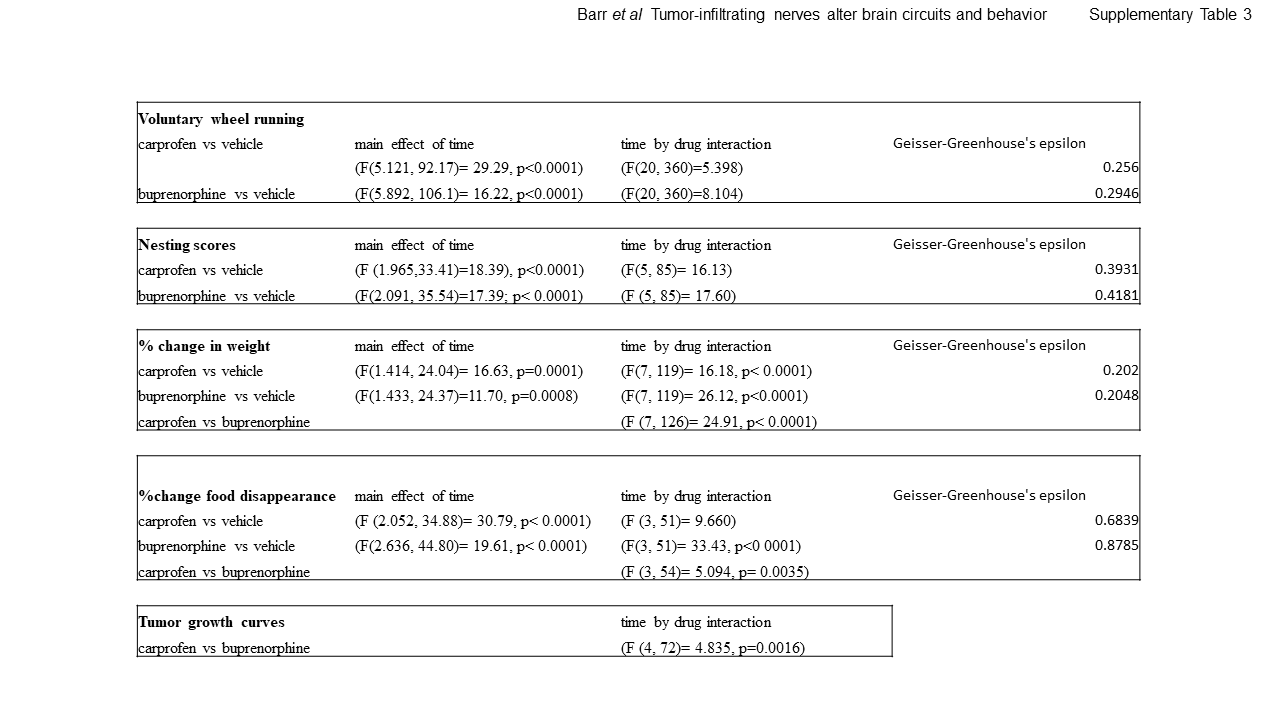

**Supplementary Table 3.** Significant effects for the statistics presented in Figure 5.

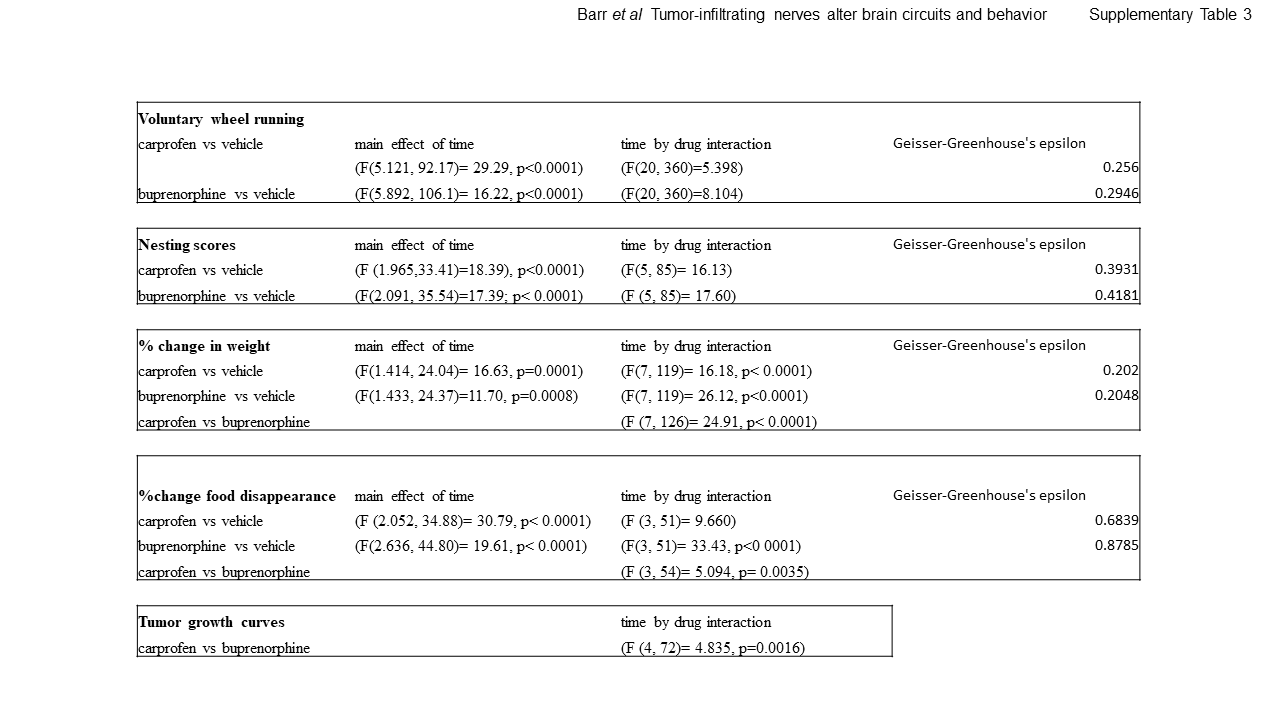

**Supplementary Table S3.** Significant effects for the statistics presented in Figure 5.
